# Ventral pallidum neurons signal relative threat

**DOI:** 10.1101/2020.06.04.135145

**Authors:** Mahsa Moaddab, Madelyn H. Ray, Michael A. McDannald

## Abstract

Ventral pallidum (VP) neurons scale firing increases to reward value and decrease firing to aversive cues. Anatomical connectivity suggests a critical role for the VP in threat-related behavior. Here we tested whether firing decreases in VP neurons conform to relative threat by recording single units while male rats discriminated cues predicting unique foot shock probabilities. Rats behavior and VP single unit firing discriminated danger, uncertainty and safety cues. We found that two VP populations (Low firing and Intermediate firing) signaled relative threat, proportionally decreased firing according shock probability: danger < uncertainty < safety. Low firing neurons showed reward firing increases, consistent with a general signal for relative value. Intermediate firing neurons were unresponsive to reward, revealing a specific signal for relative threat. The results suggest an integral role for the VP in threat-related behavior.

## Introduction

Environmental threats lie on a continuum from danger to safety, with most threats involving uncertainty. Determining relative threat – where present threat lies on the continuum – allows for an adaptive fear response. Brain regions essential to fear, most notably the central amygdala^1–3^ and basolateral amygdala (BLA)^4–8^, must be necessary to signal and utilize relative threat. At the same time, threat learning and behavior is the product of a larger neural network^9,10^ that includes regions traditionally implicated in reward^11–14^. The ventral pallidum (VP) is a compelling candidate for a neural source of relative threat. Anatomically, the VP is positioned to send and receive threat information. So although best known as an output of the mesolimbic system^15,16^, the VP receives direct projections from the central amygdala^17–20^ and has reciprocal projections with the BLA^21,22^.

The VP is consistently implicated in reward processes^20,23–28^. VP neurons acquire firing to reward-predictive cues^29–31^ and show differential firing to cues predicting different reward sizes^32,33^. VP neurons change their firing when a taste changes from palatable to aversive^34^. More recent work suggests the VP is a source of relative reward value. Single VP neurons track palatability in a multi-reward setting, showing firing increases that scale with palatability^35^. Yet, VP neurons do not exclusively signal relative reward value. The VP contributes to the formation of a conditioned taste aversion^20,36–38^ and VP neurons can acquire responding to aversive cues^39^. Most pertinent, VP neurons can show opposing changes in firing to rewarding and aversive cues. In mice, a VP population shows firing increases to rewarding cues, but firing decreases to aversive cues^18^. VP neurons showing firing increases to rewarding cues and decreases to aversive cues have also been observed in monkeys^40^. Consistent across both studies, mouse and monkey VP contained a separate population that showed firing increases to rewarding and aversive cues, indicative of salience signaling^18,40–42^.

Here we test the hypothesis that VP neurons signal relative threat, particularly through firing decreases. We recorded VP single unit activity from rats undergoing probabilistic fear discrimination consisting of cues for danger (*p*=1.00), uncertainty (*p*=0.25) and safety (*p*=0.00). Using foot shock outcome permitted direct examination of threat, as shock-predictive cues produce species specific defensive behavior^43,44^. Fear discrimination took place over a baseline of reward seeking^45^ and complete behavioral discrimination was observed. The behavior/recording approach allowed us to reveal activity patterns reflecting relative threat, relative value spanning threat and reward through opposing changes in firing, as well as salience.

## Results

Male, Long Evans rats (n = 14) were moderately food-deprived and trained to nose poke in a central port in order to receive a reward (food pellet). Nose poking was reinforced throughout fear discrimination, but poke-reward and cue-shock contingencies were independent. During fear discrimination, three distinct auditory cues predicted unique foot shock probabilities: danger (*p*=1.00), uncertainty (*p*=0.25), and safety (*p*=0.00) (Fig. 1a). Each fear discrimination session consisted of 16 trials: 4 danger, 2 uncertainty-shock, 6 uncertainty-omission, and 4 safety, mean 3.5 min inter-trial interval (ITI). Each trial started with a 20 s base-line period followed by 10 s cue presentation. Foot shock (0.5 mA, 0.5 s) was administered 2 s following cue offset on shock trials (Fig. 1b). Trial order was randomized for each rat, each session. Fear was measured by the suppression of rewarded nose poking. A suppression ratio was calculated by comparing nose poke rates during baseline and cue periods (see methods). After eight discrimination sessions, rats were implanted with drivable microelectrode bundles dorsal to the VP (Fig. 1c). Following recovery, single unit activity was recorded while rats underwent fear discrimination. The microelectrode bundle was advanced through the VP in ∼84 μm steps every other day.

**Fig 1.**
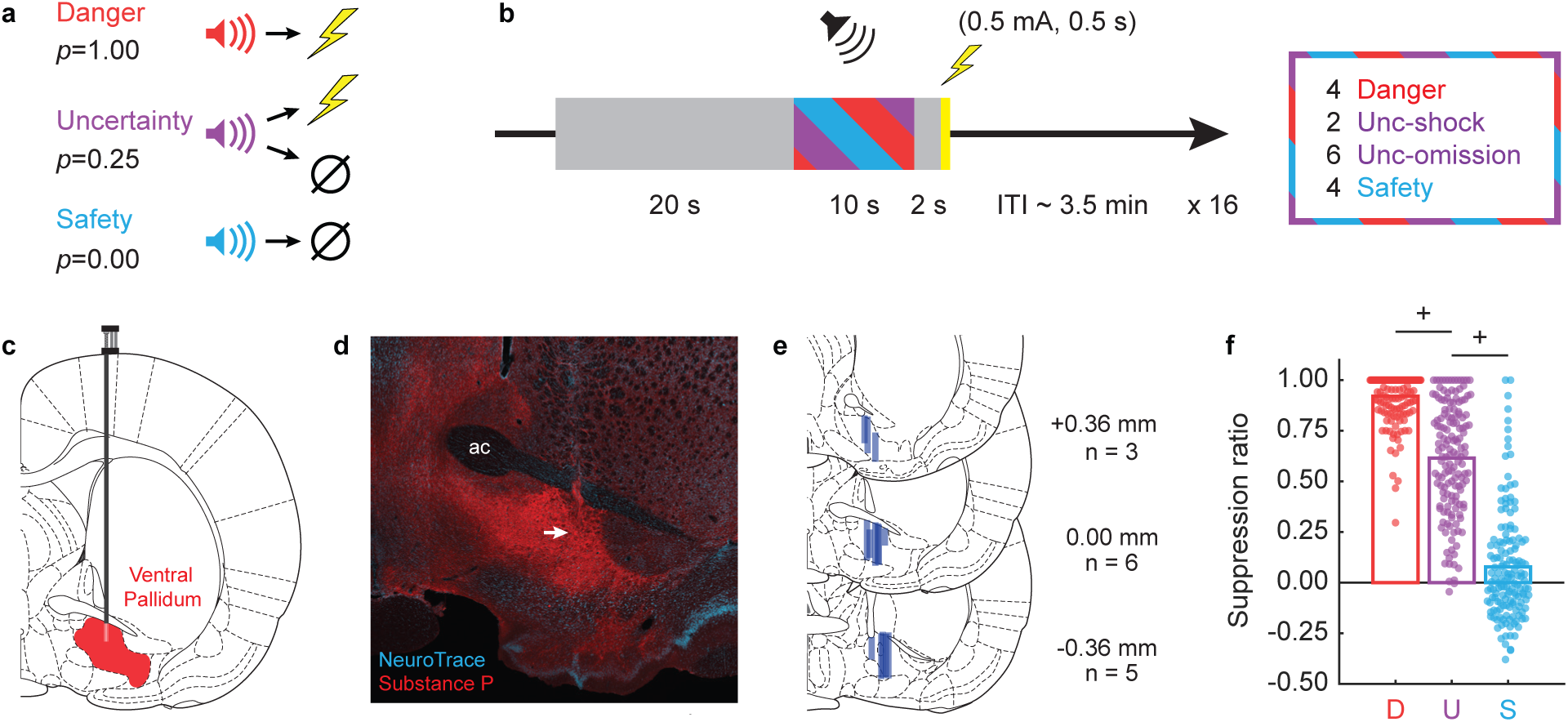
Fear discrimination, histology and behavior. (**a**) Pavlovian fear discrimination consisted of three auditory cues, each associated with a unique probability of foot shock: danger (*p*=1.00, red), uncertainty (*p*=0.25, purple) and safety (*p*=0.00, blue). (**b**) Each trial started with a 20 s baseline period followed by 10 s cue period. Foot shock (0.5 mA, 0.5 s) was administered 2 s following the cue offset in shock and uncertainty-shock trials. Each session consisted of 16 trials: four danger trials, two uncertainty-shock trials, six uncertainty-omission trials and four safety trials with an average inter-trial interval (ITI) of 3.5 min. (**c**) After training, a drivable 16 microelectrode bundle was implanted dorsal to the VP. (**d**) Example of substance P immunohistochemistry (red) showing the location of the recording site (marked by the white arrow) within the boundaries of the VP (NeuroTrace in blue). (**e**) Histological reconstruction of microelectrode bundle placements (n = 14) in the VP are represented by blue bars, bregma level indicated. (**f**) Mean (bar) and individual session (data points) suppression ratio for each cue (D, danger; U, uncertainty; S, safety) is shown for all recording sessions (n = 152) with cue-responsive neurons (n = 257). ^+^95% bootstrap confidence interval for differential suppression ratio does not contain zero.

Electrode placement was confirmed with immunohistochemistry for substance P^21^(Fig. 1d). Only placements below the anterior commissure (ac) and within the dense substance P field were accepted. A total of 435 VP neurons were recorded from 14 rats over 194 sessions (Fig.1e). To identify cue-responsive neurons, we compared baseline firing rate (mean of 10 s prior to cue onset) to firing rate during the first 1 s and last 5 s of danger, uncertainty and safety (paired, two-tailed t-test, *p*<0.05). A neuron was considered cue-responsive if it showed a significant increase or decrease in firing to any cue in either period. This screen identified 257 cue-responsive neurons (∼59% of all recorded neurons) from 152 sessions, with at least one cue-responsive neuron identified in each of the 14 rats. All remaining analyses focused on cue-responsive neurons (n = 257) and the discrimination sessions (n = 152) in which they were recorded.

Rats showed complete discrimination during sessions in which cue-responsive neurons were recorded. Suppression ratios were high to danger, intermediate to uncertainty, and low to safety (Fig. 1f). Analysis of variance (ANOVA) for suppression ratio [factor: cue (danger, uncertainty, and safety)] revealed a main effect of cue (F_2,302_ = 624.90, *p*=4.63 x 10^−108^, partial eta squared (η _p_^2^) = 0.81, observed power (op) = 1.00). Differential suppression ratios were observed for each cue pair. The 95% bootstrap confidence interval for differential suppression ratio did not contain zero for danger vs. uncertainty (mean = 0.31, 95% CI [(lower bound) 0.27, (upper bound) 0.34]) and uncertainty vs. safety (M = 0.54, 95% CI [0.48, 0.59]) (Fig. 1f). Observing robust fear discrimination permits a rigorous examination of VP threat-related responding.

### Diversity in VP baseline firing and threat responding

Plotting baseline firing rate, cue and reward firing for each neuron revealed diversity of patterned firing with three prominent features: (1) a mixture of cue-excited and cue-inhibited neurons (2) showing greatest firing changes to danger and (3) marked variation in baseline firing rate (Fig. 2). To reveal functional VP neuron-types, we averaged first 1 s and last 5 s danger firing for each neuron, designating neurons with positive values as cue-excited (n = 131, ∼51% of all cue-responsive neurons; Fig. 2 top) and neurons with negative values as cue-inhibited (n = 126, ∼49% of all cue-responsive neurons; Fig. 2 bottom). We used analysis of covariance (ANCOVA) to determine if baseline firing rate – a candidate marker for neuron-type^46^ – informed the firing pattern of cue-excited neurons. ANCOVA [covariate: baseline firing rate; within factors: cue (danger, uncertainty, and safety) and bin (250 ms bins 2 s prior to cue onset → 2 s following cue offset)] found no baseline x cue x bin interaction (F_110,14080_ = 1.12, *p*=0.18, η_p_^2^ = 0.009, op = 1.00). Because baseline firing rate did not inform the pattern of cue firing, all remaining analyses treated cue-excited neurons as a single population.

**Fig 2.**
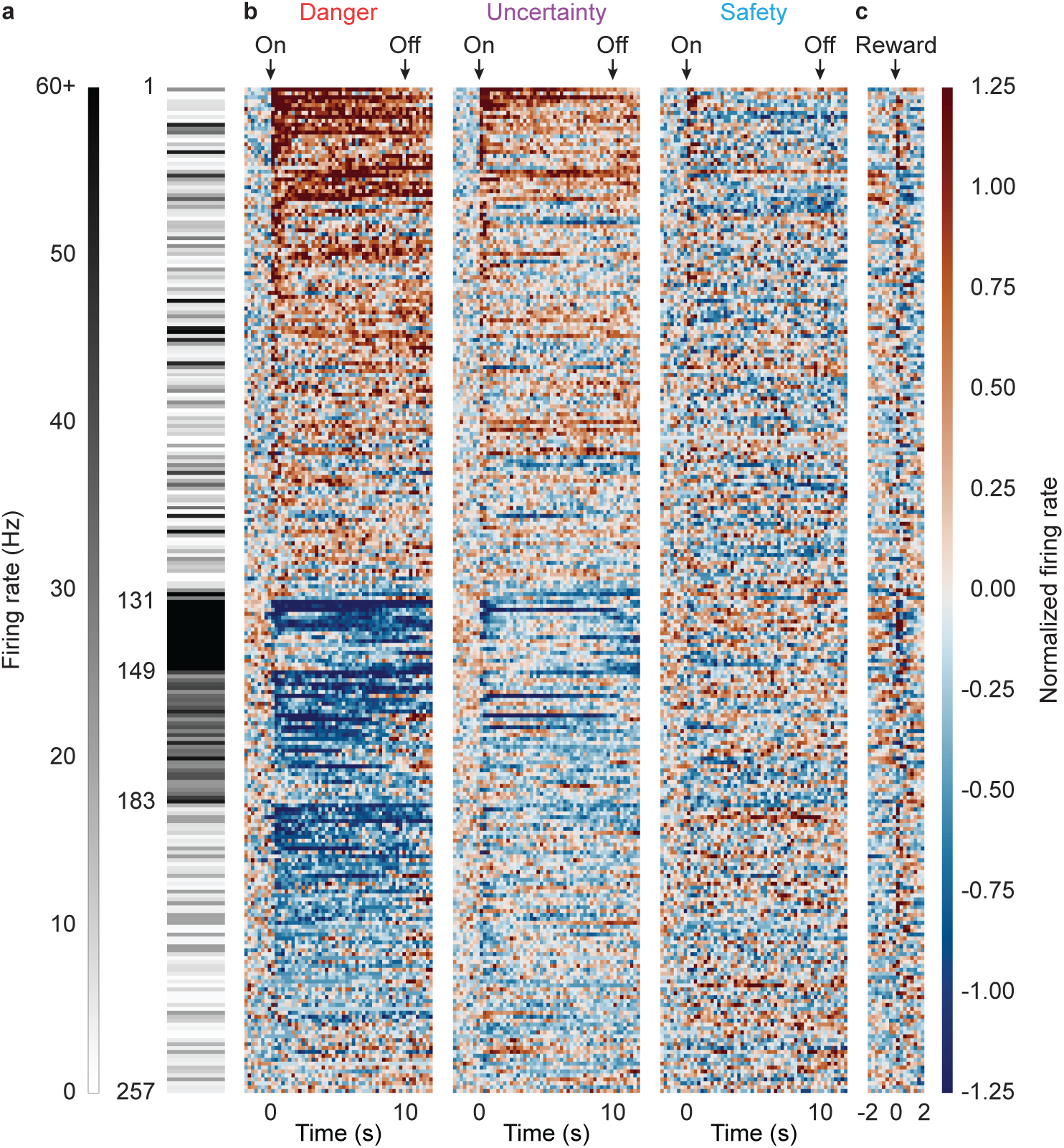
Heat plot of cue-responsive neurons. (**a**) Heat plot showing mean baseline firing rate (10 s prior to cue onset) for each cue-responsive neuron (n = 257). Color scale for baseline firing rate is shown to the left, white indicates low baseline firing rate and black high baseline firing rate. (**b**) Mean normalized firing rate for each cue-responsive neuron (n = 257), from 2 s prior to cue onset to 2 s following cue offset, in 250 ms bins for each of the three trial types: danger, uncertainty and safety. Cue onset (On) and offset (Off) are indicated by black arrows. All cue-responsive neurons are sorted by the direction of their response to danger cue (cue-excited, n = 131, maroon, top; cue-inhibited, n = 126, dark blue, bottom). Color scale for normalized firing rate is shown to the right. A normalized firing rate of zero is indicated by the color white, with greatest increases maroon and greatest decreases dark blue. (**c**) Mean normalized firing rate for each cue-responsive neuron (n = 257) from 2 s prior to 2 s following reward delivery (colors maintained from b). Reward delivery is indicated by black arrow.

We applied the same approach to determine if baseline firing rate informed responding by cue-inhibited neurons. Now, ANCOVA revealed a significant baseline x cue x bin interaction (F_110,13420_ = 1.93, *p*=2.34 × 10^−8^, η_p_^2^ = 0.02, op = 1.00). To identify distinct functional types, we used k-means clustering for baseline firing and four additional characteristics: coefficient of variance^47,48^, coefficient of skewness^48^, waveform half-duration^49^, and waveform amplitude ratio^49^ (see methods for full description of each). Cue-inhibited neurons could be divided into three clusters that differed primarily in baseline firing rate: Low firing (n = 74), Intermediate firing (n = 34), and High firing (n = 18); (firing and waveform characteristics can be found in Fig. S1). Between-cluster differences in patterned cue firing were confirmed by ANOVA returning a significant cluster x cue x bin interaction for all comparisons (Low vs. Intermediate, Low vs. High, and Intermediate vs. High; all F > 1.40, all *p*<0.005). The remaining cue-inhibited analyses focus on Low, Intermediate and High firing neurons.

### Differential inhibition of firing is maximal to danger

If VP cue-inhibited neurons signal relative threat through firing decreases, greatest firing inhibition should be observed to danger, the cue associated with the highest shock probability. Lesser and more similar inhibition of firing should be observed to uncertainty and safety; whose foot shock probabilities are closer to one another. To determine if differential firing was observed, we separately performed ANOVA for Low, Intermediate and High firing neurons [factors: cue (danger, uncertainty, and safety) and bin (250 ms bins from 2 s prior to cue onset → 2 s following cue offset)]. The cue response pattern for Low firing neurons complied with requirements of a neural signal for relative threat. Low firing neurons showed greatest inhibition to danger, modest inhibition to uncertainty and no inhibition to safety. The relative firing pattern was maintained throughout cue presentation (Fig. 3a). Confirming differential firing, ANOVA for normalized firing rate (Z-score) for the 74 Low firing neurons revealed a significant main effect of cue (F_2,142_ = 23.45, *p*=1.58 × 10^−9^, η _p_^2^ = 0.25, op = 1.00), bin (F_55,3905_ = 4.66, *p*=1.18 × 10^−26^, η _p_^2^ = 0.06, op = 1.00), and most critically a significant cue x bin interaction (F_110,7810_ = 3.27, *p*=1.80 × 10^−27^, η _p_^2^ = 0.04, op = 1.00).

**Fig 3.**
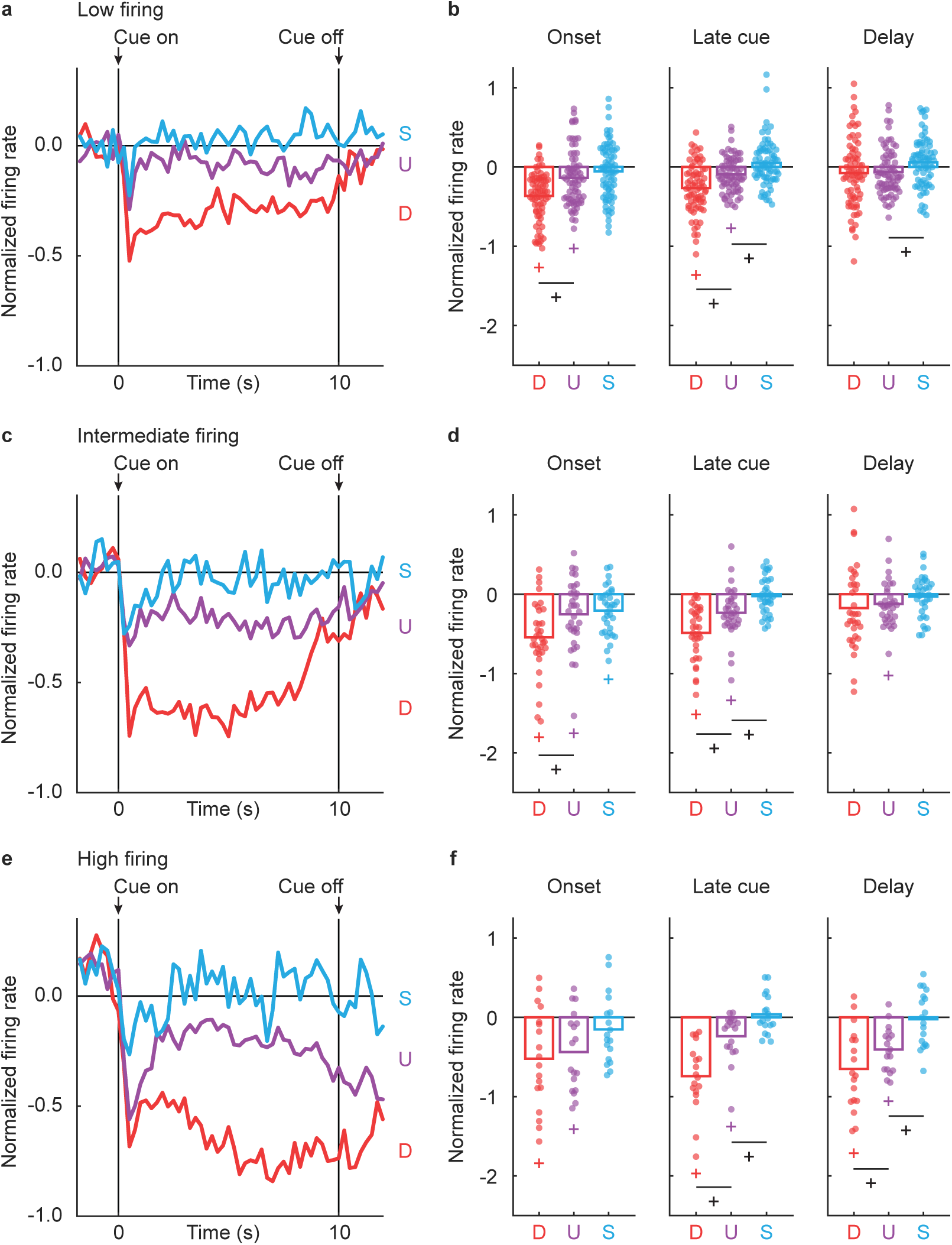
Differential firing in cue-inhibited population. (**a**) Mean normalized firing rate to danger (red), uncertainty (purple) and safety (blue) is shown from 2 s prior to cue onset to 2 s following cue offset for the Low firing neurons (n = 74). Cue onset and offset are indicated by vertical black lines. (**b**) Mean (bar) and individual (data points), normalized firing rate for Low firing neurons (n = 74) during the first 1 s cue interval (onset, left), the last 5 s cue interval (late cue, middle), and 2 s following cue offset (delay, right) are shown for each cue (D, danger; U, uncertainty; and S, safety). (**c**) Mean normalized firing rate for the Intermediate firing neurons (n = 34), shown as in a. (**d**) Mean (bar) and individual (data points), normalized firing rate for Intermediate firing neurons (n = 34), as in b. (**e**) Mean normalized firing rate for the High firing neurons (n = 18), shown as in a. (**f**) Mean (bar) and individual (data points), normalized firing rate for High firing neurons (n = 18), as in b. ^+^95% bootstrap confidence interval for differential cue firing does not contain zero (black plus signs). ^+^95% bootstrap confidence interval for normalized firing rate does not contain zero (colored plus signs).

To determine if population-level patterns were observed in single units, we constructed 95% boot strap confidence intervals for normalized firing rate for each cue (compared to zero), as well as for differential firing: (danger vs. uncertainty) and (uncertainty vs. safety). Separate 95% bootstrap confidence intervals were constructed for cue onset (first 1 s cue interval), late cue (last 5 s cue interval), and delay (2 s following cue offset) periods. Observing 95% bootstrap confidence that did not contain zero supports the interpretation that differential firing was observed.

Low firing neurons showed selective inhibition of firing to threat cues at onset (danger: M = −0.37, 95% CI [-0.44, −0.29]; uncertainty: M = −0.14, 95% CI [-0.22, −0.06]) and during late cue (danger: M = −0.27, 95% CI [-0.34, −0.19]; uncertainty: M = −0.09, 95% CI [-0.15, −0.04]) (Fig. 3b, colored plus signs). Low firing neurons showed differential firing to danger and uncertainty at onset (M = −0.23, 95% CI [-0.31, −0.13]; Fig. 3b, left) and during late cue (M = −0.17, 95% CI [-0.25, −0.09]; Fig.3b, middle). Similar firing was observed to uncertainty and safety at onset (M = −0.08, 95% CI [-0.19, 0.02]; Fig. 3b, left), but differential firing was observed during late cue (M = −0.14, 95% CI [-0.25, −0.04]; Fig. 3b, middle) and delay (M = −0.12, 95% CI [-0.26, −6.32 × 10^−4^]; Fig. 3b, right).

Intermediate firing neurons (n = 34) also showed patterned firing consistent with a neural signal for relative threat. At cue onset, there was greatest firing inhibition to danger, lesser inhibition to uncertainty and least inhibition to safety. Inhibition of firing that was specific to danger and uncertainty was maintained for the remainder of cue presentation (Fig. 3c). In support, ANOVA revealed a significant main effect of cue (F_2,66_ = 27.25, *p*=2.36 × 10^−9^, η _p_^2^ = 0.45, op = 1.00), bin (F _55,1815_ = 7.62, *p*=1.52 × 10^−50^, η _p_^2^ = 0.19, op = 1.00), and a significant cue x bin interaction (F_110,3630_ = 3.90, *p*=2.20 × 10^−36^, η_p_ ^2^ = 0.11, op = 1.00). Single unit analysep confirmed the ANOVA results. Intermediate firing neurons were inhibited to all cues at onset (danger: M = −0.54, 95% CI [-0.69, −0.39]; uncertainty: M = −0.25, 95% CI [-0.40, −0.08]; safety: M = −0.21, 95% CI [-0.30, −0.11]), but were selectively inhibited to danger and uncertainty during late cue (danger: M = −0.49, 95% CI [-0.60, −0.36]; uncertainty: M = −0.23, 95% CI [-0.34, −0.12]) (Fig. 3d, colored plus signs). Differential inhibition of firing to danger and uncertainty was observed at cue onset (M = −0.29, 95% CI [-0.44, −0.14]; Fig. 3d, left), and during late cue (M = −0.26, 95% CI [-0.42, −0.10]; Fig. 3d, middle). Like for Low firing neurons, differential inhibition of firing was not observed to uncertainty and safety at cue onset (M = −0.05, 95% CI [-0.23, 0.14]; Fig. 3d, left), but was observed during late cue (M = −0.21, 95% CI [-0.34, −0.05]; Fig. 3d, middle).

High firing neurons (n = 18) showed a distinct firing pattern. Inhibition of firing was observed to danger and uncertainty at onset, yet differential firing to each cue pair was not observed. Firing decreases specific to danger and uncertainty continued through the late cue and delay period, only now full discrimination observed in each period (Fig. 3e). ANOVA revealed main effects of cue (F_2,34_ = 19.90, *p*=2.00 × 10^−6^, η_p_^2^ = 0.54, op = 1.00), and bin (F_55,935_ = 7.84, *p*=3.82 × 10^−47^, η_p_^2^ = 0.32, op = 1.00), as well as a cue x bin interaction (F_110,1870_ = 3.11, *p*=7.56 × 10^−23^, η_p_^2^ = 0.16, op = 1.00). High firing neurons were inhibited to danger and uncertainty during all periods: onset (danger: M = −0.52, 95% CI [-0.84, −0.21]; uncertainty: M = −0.44, 95% CI [-0.69, −0.20]), late cue (danger: M = −0.74, 95% CI [-0.93, −0.49]; uncertainty: M = −0.23, 95% CI [-0.35, −0.009]), and delay (danger: M = −0.65, 95% CI [-0.92, −0.42]; uncertainty: M = −0.40, 95% CI [-0.55, −0.27]) (Fig. 3f, colored plus signs). Differential firing was not observed at onset (danger vs. uncertainty: M = −0.09, 95% CI [-0.27, 0.07]; uncertainty vs. safety: M = −0.28, 95% CI [-0.62, 0.001]; Fig. 3f, left), but was subsequently observed for all cue pairs (danger vs. uncertainty, late cue: M = −0.51, 95% CI [-0.73, −0.22]; delay: M = −0.24, 95% CI [-0.42, −0.02]), (uncertainty vs. safety, late cue: M = −0.28, 95% CI [-0.44, −0.08]; delay: M = −0.39, 95% CI [-0.64, −0.14]) (Fig. 3f).

Population and single unit firing analyses reveal Low, Intermediate and High firing neurons are candidate sources of relative threat signaling. Even more, positive firing relationships were commonly observed for threat cues, danger and uncertainty, but zero or even negative firing relationships were observed for uncertainty and safety (Fig. S2). Inhibition of firing was not simply due to the cessation of nose poking. Pauses in nose poking in the absence of cues during the inter-trial interval were insufficient to inhibit the activity of Low, Intermediate and High firing neurons (Fig. S3). Of course, differential cue firing would also be expected of a neural signal for fear output. Given that our rats showed complete behavioral discrimination of danger, uncertainty and safety; inhibition of VP firing could reflect fear output, rather than relative threat.

### Low and Intermediate firing neurons signal relative threat

We used single unit linear regression to determine the degree to which VP single unit activity reflected fear output and relative threat (see methods). Fear output and relative threat could be dissociated because rats showed greater fear to uncertainty (D > U >> S) than would be expected based on its foot shock probability (D >>> U > S; Fig. 1f). For each single unit, we calculated the normalized firing rate for each trial (16 total: 4 danger, 8 uncertainty, and 4 safety trials) for a total of 14 s (1 s bins; 2 s prior to cue onset, 10 s cue presentation, and 2 s following cue offset). The fear output regressor was the cue suppression ratio for that specific trial. The relative threat regressor was the foot shock probability assigned to each cue (danger = 1.00, uncertainty = 0.25, and safety = 0.00). Regression output was a beta coefficient for each regressor, quantifying the strength (greater distance from 0 = stronger) and direction (>0 = positive and <0 = negative) of the predictive relationship between each regressor and single unit firing. Regression allowed us to determine whether the firing of each VP neuron was better described by the rat’s behavior in that session (fear output) or the shock probability associated with each cue (relative threat)^50,51^.

Low firing neurons (n = 74) signaled relative threat (Fig. 4a). Neither fear output nor relative threat signaling was observed prior to cue onset. Relative threat signaling – decreases in firing that scaled to shock probability – was apparent immediately upon cue onset and was maintained for the entirety of cue presentation (Fig. 4a, green plus signs). Fear output signaling only emerged in the final two seconds of cue presentation (Fig. 4a, gray plus signs). ANOVA for beta coefficient [factors: regressor (relative threat and fear output) and interval (1 s bins from 2 s prior to cue onset → 2 s following cue offset)] found main effect of regressor (F_1,72_ = 4.38, *p*=0.04, η _p_^2^ = 0.06, op = 0.54), but most importantly a significant regressor x interval interaction (F_13,936_ = 3.49, *p*=2.5 × 10^−5^, η _p_^2^ = 0.05, op = 1.00). The mean beta coefficient across cue presentation was below zero for relative threat (M = −0.38, 95% CI [-0.52, −0.24]), but not fear output (M = −0.06, 95% CI [-0.17, 0.04]) (Fig. 4b, left, green plus sign). Moreover, the mean beta coefficient was more negative for relative threat than for fear output (M = −0.32, 95% CI [-0.52, −0.11]; Fig. 4b, left), but not during the delay period (M = 0.12, 95% CI [-0.13, 0.38]; Fig. 4b, right).

**Fig 4.**
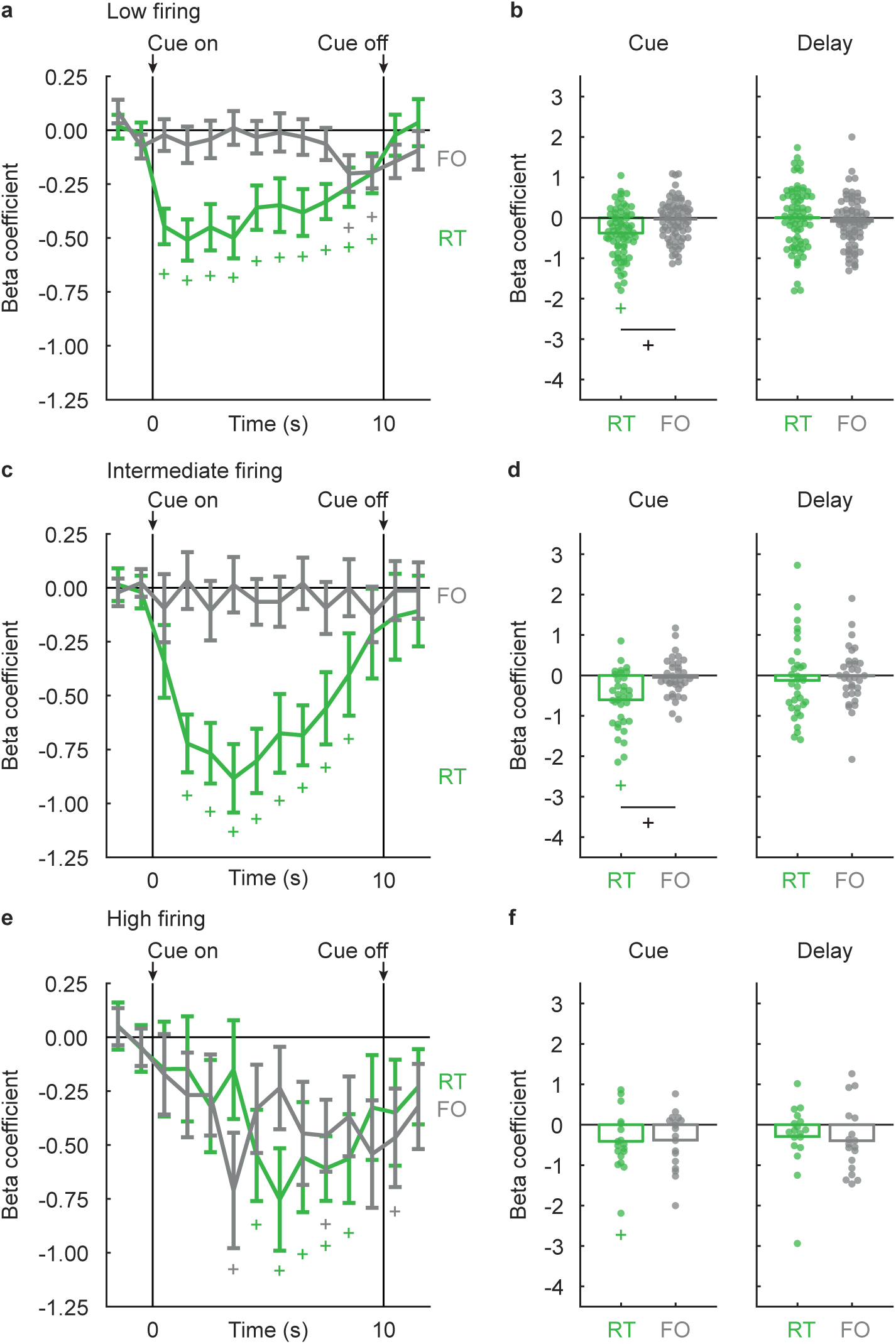
Low firing and Intermediate firing neurons signal relative threat. (**a**) Mean ± SEM beta coefficients are shown for each regressor (RT, relative threat, green; FO, fear output, gray), from 2 s prior to cue onset to 2 s following cue offset in 1 s intervals, for the Low firing neurons (n = 74). Cue onset and offset are indicated by vertical black lines. (**b**) Mean (bar) and individual (data points), beta coefficient for each regressor (RT, relative threat, green; FO, fear output, gray) during the entire 10 s cue presentation (cue), and 2 s following cue offset (delay) are shown for Low firing neurons (n = 74). (**c**) Mean ± SEM beta coefficients for the Intermediate firing neurons (n = 34), shown as in a. (**d**) Mean (bar) and individual (data points), beta coefficients for Intermediate firing neurons (n = 34), as in b. (**e**) Mean ± SEM beta coefficients for the High firing cluster (n = 18), shown as in a. (**f**) Mean (bar) and individual (data points), beta coefficients for High firing neurons (n = 18), as in b. ^+^95% bootstrap confidence interval for differential beta coefficient does not contain zero (black plus signs). ^+^95% bootstrap confidence interval for beta coefficient does not contain zero (colored plus signs).

Intermediate firing neurons exclusively signaled relative threat (Fig. 4c). Neither fear output nor relative threat signaling were observed prior to cue onset. Relative threat signaling emerged 1 s after cue onset and was maintained for all cue intervals except the last (Fig. 4c, green plus signs). Fear output signaling was not observed in any interval (Fig. 4c). In support, ANOVA revealed main effect of regressor (F_1,32_ =, *p*=0.015, η _p_^2^ = 0.17, op = 0.70), as well as a significant regressor x interval interaction (F_13,416_ = 2.41, *p*=0.004, η _p_^2^ = 0.07, op = 0.98). Mean beta coefficient for relative threat (M = −0.60, 95% CI [-0.83, −0.33]), but not fear output (M = −0.05, 95% CI [-0.24, 0.13]), were below zero across cue presentation (Fig. 4d, left, green plus sign). Further, the mean beta coefficient was more negative for relative threat than for fear output during the cue period (M = −0.56, 95% CI [-0.96, −0.13]; Fig.4d, left), but not during the delay period (M = −0.10, 95% CI [-0.70, 0.35]; Fig. 4d, right).

High firing neurons (n = 18) showed a unique pattern, with weaker signals for relative threat and fear output only emerging during late cue (Fig. 4e). ANOVA found only a main effect of interval (F_13,221_ = 7.28, *p*=8.23 × 10^−12^, η _p_^2^ = 0.30, op = 1.00). The mean beta coefficient across cue presentation was below zero for relative threat (M = −0.41, 95% CI [-0.77, −0.06]), but not fear output (M = −0.37, 95% CI [-0.68, 0.02]) (Fig. 4f, left, green plus sign). However, relative threat and fear output beta coefficients did not differ from one another during the cue period (M = −0.02, 95% CI [-0.76, 0.51]; Fig. 4f, left).

### Low firing neurons show opposing responses to threat and reward

Regression revealed that VP neurons signal relative threat through firing decreases. While our behavioral procedure is optimized to examine threat-related firing, measuring fear with conditioned suppression permitted us to record neural activity around reward delivery. Although not explicitly cued through the speaker, each reward delivery was preceded by a brief sound caused by the advance of the pellet feeder. We asked if reward-related firing was observed in Low, Intermediate and High firing neurons. Increases in reward fir ing – opposing the direction to threat – would indicate relative value signaling that spans reward and threat. The absence of reward firing would indicate specific signaling of relative threat. Decreases in firing would indicate salience signaling.

To determine reward-related firing, and possible differences between Low, Intermediate and High firing neurons, we performed repeated measures ANOVA for normalized firing rate [factors: cluster (Low vs. Intermediate vs. High) and bin (16 total: 250 ms bins from 2 s prior → 2 s following reward delivery)]. Low firing neurons sharply increased responding following reward delivery, and this firing increase was absent in Intermediate and High firing neurons (Fig. 5a). In support, ANOVA found a cluster x bin interaction (F_30,1815_ = 3.84, *p*=1.72 × 10^−11^, η _p_^2^ = 0.06, op = 1.00). Performing separate ANOVA for each cluster revealed a main effect of bin in Low firing neurons (F_15,1065_ = 8.07, *p*=1.51 × 10^−17^, η _p_^2^ = 0.10, op = 1.00), but not Intermediate (F_15,495_ = 1.21, *p*=0.26, η _p_^2^ = 0.04, op = 0.77) and High firing neurons (F = 1.06, *p*=0.40, η _p_^2^ = 0.06, op = 0.68). Pre-reward responding by Low firing neurons hovered around zero (M = −0.06, 95% CI [-0.15, 0.01]), while post-reward firing exceeded pre-reward firing (M = 0.34, 95% CI [0.15, 0.53]) and differed from zero (M = 0.28, 95% CI [0.14, 0.42]) (Fig. 5b, black plus signs). By contrast, pre- and post-reward firing never differed from zero for Intermediate (pre-reward: M = 2.60 × 10^−4^, 95% CI [-0.17, 0.15]; post-reward: M = −0.14, 95% CI [-0.41, 0.12]; Fig. 5c) and High firing neurons (pre-reward: M = 0.02, 95% CI [-0.21, 0.27]; post-reward: M = −0.29, 95% CI [-0.80, 0.19]; Fig. 5d).

**Fig 5.**
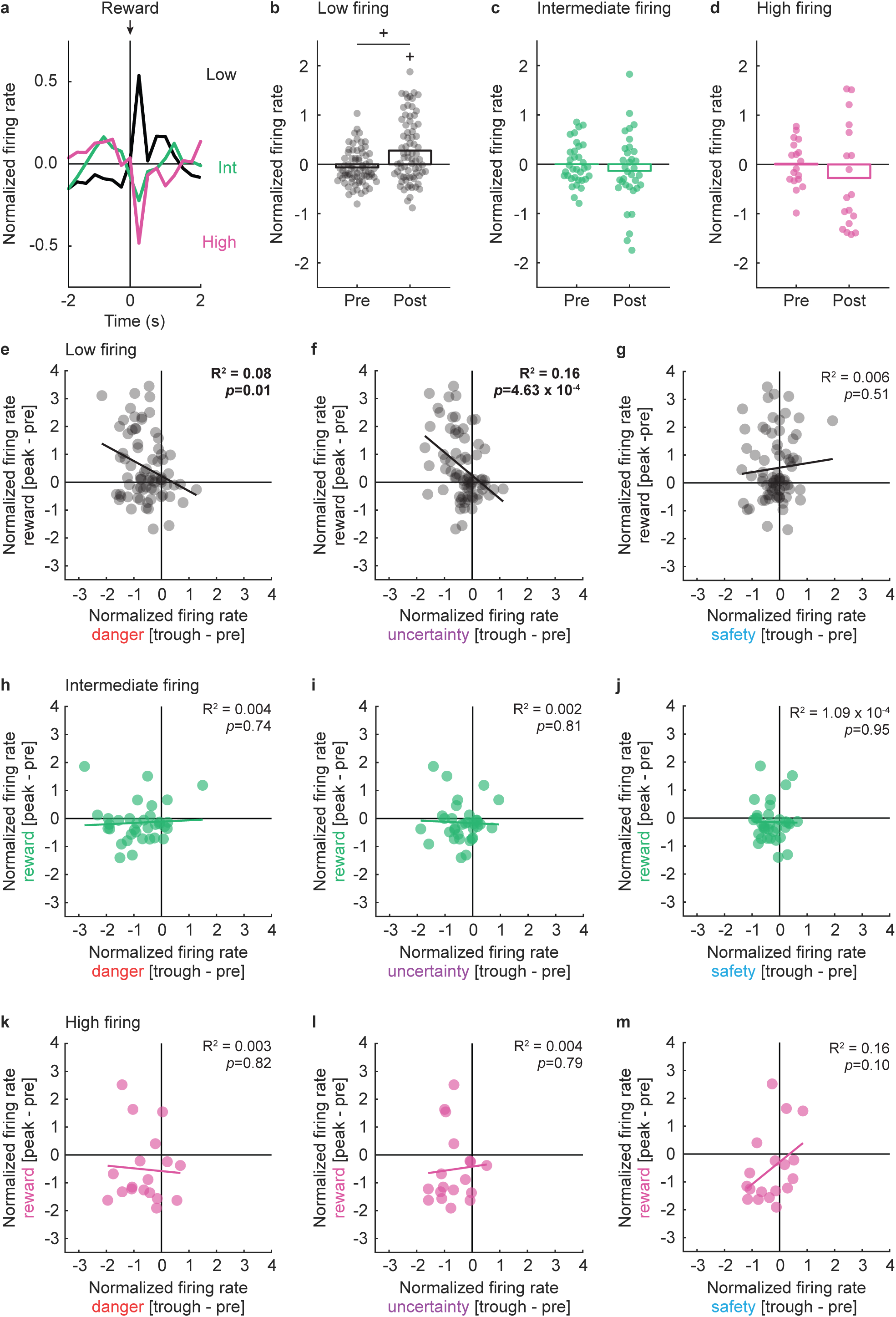
Low firing neurons show opposing responses to threat and reward. (**a**) Mean normalized firing rate to reward is shown 2 s prior to and 2 s after reward delivery for the Low (n = 74, black), Intermediate (n = 34, turquoise), and High (n = 18, pink) firing neurons. Reward delivery is indicated by black arrow. (**b**) Mean (bar) and individual (data points), normalized firing rate for Low firing neurons (n = 74, black) are shown during 500 ms interval prior (pre) to and 500 ms interval after (post) reward delivery. (**c** and **d**) Mean and individual, normalized firing rate for (**c**) Intermediate (n = 34, turquoise), and (**d**) High (n = 18, pink) firing neurons, as in b. (**e**) Mean normalized firing rate to reward (250 ms prior to reward delivery to 250 ms following reward delivery, [peak - pre]) vs. danger (the second 250 ms of cue, [trough - pre], red) is plotted for Low firing neurons (n = 74, black). (**f** and **g**) Mean normalized firing to (**f**) reward vs. uncertainty (purple) and (**g**) reward vs. safety (blue) is plotted for Low firing neurons (n = 74, black), as in e. Trendline, the square of the Pearson correlation coefficient (R^2^) and associated p value (*p*) are shown for each graph. (**h**-**j**) Mean normalized firing rate to (**h**) reward vs. danger, (**i**) reward vs. uncertainty (**j**) reward vs. safety for Intermediate firing neurons (n = 34, turquoise), as in e-g. (**k**-**m**) Mean normalized firing rate to (**k**) reward vs. danger, (**l**) reward vs. uncertainty (**m**) reward vs. safety for High firing neurons (n = 18, pink), as in e-g. ^+^95% bootstrap confidence interval for differential reward firing does not contain zero. ^+^95% bootstrap confidence interval for normalized firing rate does not contain zero.

Not only did Low firing neurons show opposing changes in firing to threat and reward, but the magnitude of firing change was negatively correlated. Reward and danger firing were negatively correlated. That is, Low firing neurons showing greater reward firing increases, showed greater danger firing decreases (R^2^ = 0.08, *p*=0.01; Fig. 5e). Reward and uncertainty firing were also negatively correlated (R^2^ = 0.16, *p*=4.63 × 10^−4^; Fig. 5f), but zero relationship was observed for reward and safety firing (R^2^ = 0.006, *p*=0.51; Fig. 5g). Even more, cue-reward firing relationships were specific to threat. Equivalent danger-reward and uncertainty-reward correlations were observed in Low firing neurons (Fisher r-to-z-transformation, Z = 0.74, *p*=0.46), but uncertainty-reward and safety-reward correlations significantly differed (Z = 2.96, *p*=0.0031). No cue-reward firing relationships were observed for Intermediate firing (Fig. 5h-j) and High firing neurons (Fig. 5k-m), and these correlations did not differ from one another (all Z < 1, all *p*>0.3). Altogether, the results reveal VP signaling of relative threat through inhibition of cue firing. Low firing neurons signal relative threat with opposing responses to reward. Intermediate firing neurons specifically signal relative threat. High firing neurons more weakly signal a mix of fear output and relative threat.

### Differential increases in firing are maximal to danger

We identified 131 neurons (∼51% of all cue-responsive neurons) showing firing increases to danger. Cue-excited neurons sharply increased activity at onset, with greatest firing to danger, lesser to uncertainty and least to safety. Differential firing continued during the remainder of the cue and through the 2 s delay period (Fig. 6a). ANOVA for normalized firing rate [factors: cue (danger, uncertainty, and safety) and bin (250 ms bins 2 s prior to cue onset → 2 s following cue offset)] revealed main effects of cue (F_2,258_ = 68.22, *p*=1.65 × 10_^-24^_, η _p_^2^ = 0.35, op = 1.00) and bin (F_55,7095_ = 15.06, *p*=5.42 × 10^−130^, η _p_^2^ = 0.11, op = 1.00) and a cue x bin interaction (F110,14190 = 4.52, *p*=2.31 _p_ 10^−49^, η _p_^2^ = 0.03, op = 1.00).

**Fig 6.**
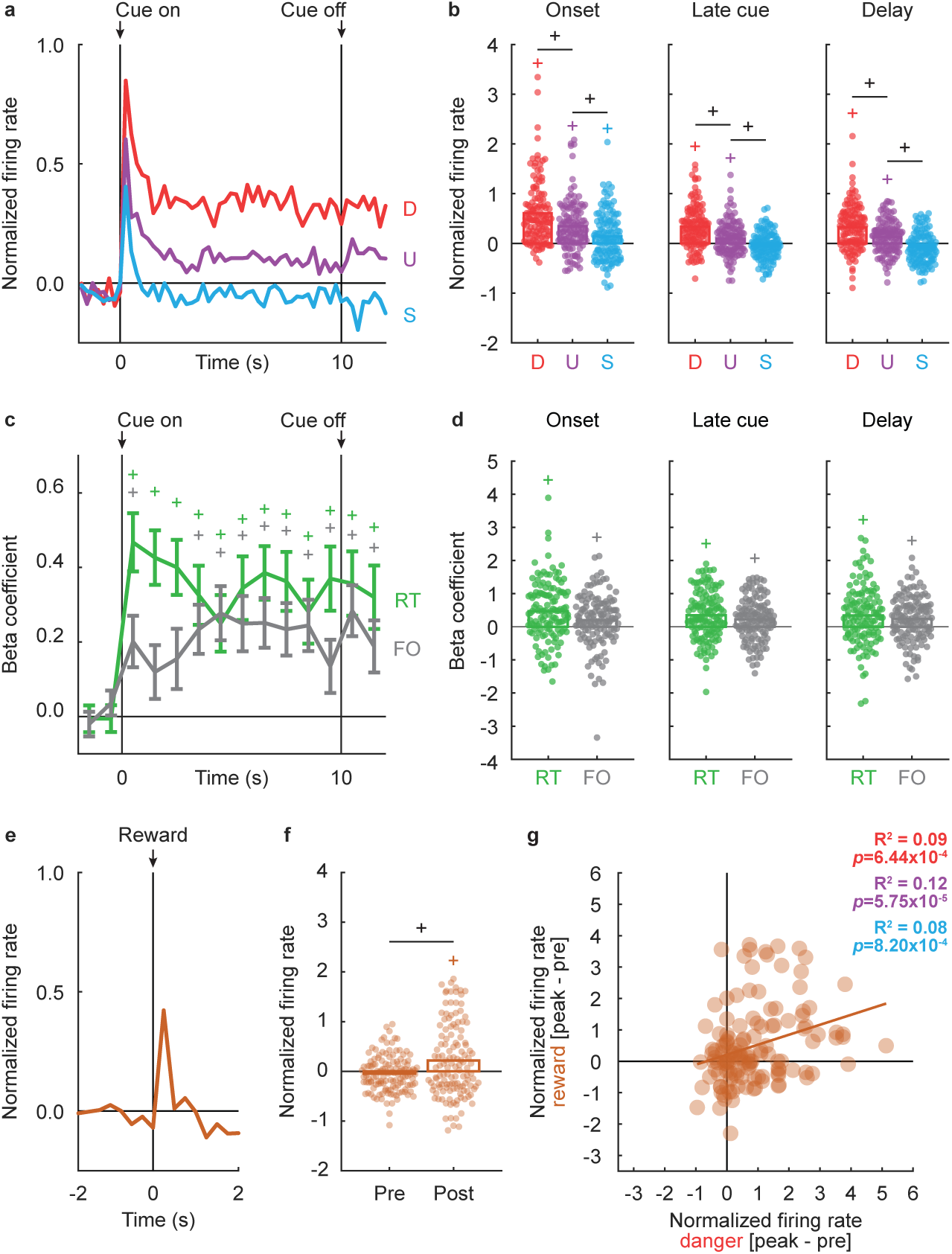
Differential cue firing by cue-excited neurons reflects relative threat and fear output. (**a**) Mean normalized firing rate to danger (red), uncertainty (purple) and safety (blue) is shown from 2 s prior to cue onset to 2 s following cue offset for the cue-excited population (n = 131). Cue onset and offset are indicated by vertical black lines. (**b**) Mean (bar) and individual (data points), normalized firing rate for cue-excited neurons (n = 131) during the first, 1 s cue interval (onset, left), the last, 5 s cue interval (late cue, middle), and 2 s following cue offset (delay, right) are shown for each cue (D, danger; U, uncertainty; S, safety). ^+^95% bootstrap confidence interval for differential cue firing rate does not contain zero (black plus signs). ^+^95% bootstrap confidence interval for normalized firing rate does not contain zero (colored plus signs). (**c**) Mean ± SEM beta coefficients are shown for each regressor (RT, relative threat, green; FO, fear output, gray), from 2 s prior to cue onset to 2 s following cue offset in 1 s intervals, for the cue-excited population (n = 131). Cue onset and offset are indicated by vertical black lines. (**d**) Mean (bar) and individual (data points), beta coefficient for each regressor (RT, relative threat, green; FO, fear output, gray) for cue-excited neurons (n = 131). ^+^95% bootstrap confidence interval for beta coefficient does not contain zero (colored plus signs). (**e**) Mean normalized firing rate to reward is shown 2 s prior to and 2 s after the reward delivery for the cue-excited population (n = 131, maroon). Reward delivery is indicated by vertical black line. (**f**) Mean (bar) and individual (data points), normalized firing rate for cue-excited neurons (n = 131, maroon) are shown during 500 ms interval prior (pre) to and 500 ms interval after (post) the reward delivery. ^+^95% bootstrap confidence interval for differential firing rate does not contain zero (black plus sign). ^+^95% bootstrap confidence interval for firing rate does not contain zero (maroon plus sign). (**g**) Mean normalized firing rate to reward (250 ms prior to reward delivery to 250 ms following reward delivery, [peak - pre]) vs. danger (the first 250 ms of cue, [peak - pre], red) is plotted for cue-excited neurons (n = 131, maroon). Trendline, the square of the Pearson correlation coefficient (R^2^) and associated p value (*p*) are shown.

Population-level firing patterns were observed in single units. Firing increases were observed to all cues at onset, but only to the threat cues, danger and uncertainty, in the remaining periods (Fig. 6b, colored plus signs). Furthermore, differential firing was observed to every cue pair in every period: danger vs. uncertainty (onset: M = 0.26, 95% CI [0.18, 0.33], late cue: M = 0.24, 95% CI [0.17, 0.30], and delay: M = 0.19, 95% CI [0.13, 0.26]), and uncertainty vs. safety (onset: M = 0.20, 95% CI [0.11, 0.30], late cue: M = 0.14, 95% CI [0.07, 0.22], and delay: M = 0.22, 95% CI [0.14, 0.31]) (Fig. 6b). Danger and uncertainty firing were positively correlated for all periods (Fig. S4a-c). By contrast, positively correlated firing to uncertainty and safety at cue onset gave way to zero correlation during late cue and negatively correlated during the delay (Fig. S4d-f). Cessation of nose poking in the absence of cues was insufficient to increase firing (Fig.S5). The pattern of differential cue firing is consistent with signaling of relative threat.

### Cue-excited neurons signal relative threat and fear output

Of course, descriptive firing analyses cannot distinguish between relative threat and fear output signaling. To do this we performed single unit, linear regression (described above). Regression revealed that single unit activity was captured by a mixture of relative threat and fear output (Fig. 6c). Cue-excited neurons showed positive beta coefficients for both regressors from cue onset through the 2 s delay period (Fig. 6c). ANOVA for beta coefficient [factors: regressor (relative threat and fear output) and interval (1 s bins from 2 s prior to cue onset → 2 s following cue offset)] found only a main effect of regressor (F_13,1690_ = 15.24, *p*=4.06 × 10^−33^, η _p_^2^ = 0.11, op = 1.00). Beta coefficients exceeding zero were observed for relative threat and fear output in nearly every interval starting with cue onset (Fig. 6c, colored plus signs). Relative threat and fear output beta coefficients did not differ from one another during any period (onset: M = 0.26, 95% CI [-0.009, 0.52], left, late cue: M = 0.12, 95% CI [-0.09, 0.35], middle, and delay: M = 0.10, 95% CI [-0.12, 0.33], right; Fig. 6d).

### Cue-excited neurons increase firing to reward

To specify if cue-excited neurons signal relative value, relative threat or salience, we examined firing around reward delivery. Reward firing decreases would support relative value signaling, no change in firing to reward would support relative threat signaling and reward firing increases would support salience signaling. Cue-excited neurons increased firing to reward (Fig. 6e). Repeated measures ANOVA for normalized firing rate [factor: bin (250 ms bins from 2 s prior → 2 s following reward delivery)] revealed a main effect of bin (F_15,1920_ = 7.24, *p*=9.32 × 10^−16^, η_p_^2^ = 0.05, op = 1.00). Single unit firing prior to reward delivery hovered around zero. Firing increases following reward delivery exceeded zero (M = 0.22, 95% CI [0.07, 0.35]; Fig. 6f, maroon plus sign), and also exceeded pre-reward firing (M = 0.26, 95% CI [0.10, 0.41]; Fig. 6f). Further supporting salience signaling, the magnitude of firing increase to each cue and reward was positively correlated. Single units showing greater firing increases at reward peak showed greater firing increases at danger (R^2^ = 0.09, *p*=6.44 × 10^−4^), uncertainty (R^2^ = 0.12, *p*=5.75 × 10^−5^) and safety peak (R^2^ = 0.08, *p*=8.20 × 10^−4^) (Fig. 6g). Finally, peak danger firing (M = 0.40, 95% CI [0.13, 0.65]), but not peak uncertainty (M = 0.13, 95% CI [-0.07, 0.35]), and safety firing (M = −0.08, 95% CI [-0.30, 0.13]), differed from peak reward firing. The results reveal a signal for relative threat within a more general signal for salience.

## Discussion

We recorded VP single unit activity while rats discriminated danger, uncertainty and safety. Revealing wide-spread threat responding, most VP neurons were maximally responsive to danger. Two cue-inhibited neuron types (Low firing and Intermediate firing) signaled relative threat, decreasing cue firing in proportion to foot shock probability (danger < uncertainty < safety). Low firing neurons increased firing following reward delivery, marking these neurons a possible source of relative value that spans threat and reward. Intermediate firing neurons more exclusively signaled relative threat, while a smaller group of High firing neurons signaled fear output and relative threat through firing decreases. Consistent with salience signaling, neurons showing firing increases to cues also increased firing to reward; cue firing reflected relative threat and fear output.

Before discussing our results, some considerations must be raised. The first concerns neurotransmitter identity. The VP is heterogeneous, containing GABAergic, glutamatergic, and cholinergic neurons^21^. We can make inferences about the neurotransmitter identity of each functional type based on firing properties and published studies. However, the present results cannot definitively tie functional populations to neurotransmitter identity. Another consideration is that our behavioral design did not manipulate reward with the same nuance as threat. This was intentional, as our goal was to examine relative threat behavior, firing and signaling. Nevertheless, our design prevents a definitive demonstration of relative value signaling that spans threat and reward. Such a demonstration would require a discrimination procedure in which 5+ cues predict unique shock and reward probabilities, observing full behavioral discrimination. We limited this study to males, in part to enable comparison of VP responding to prior reports, which have been mostly in males^27,29,32–35,37,38,42,52–54^. So far, no differences in VP activity/function have been found in studies that examined biological sex^18,39,55^. Our laboratory has observed complete and comparable fear discrimination in male and female rats^11,51,56,57^. We predict that equivalent relative threat signaling will be observed in female and male VP neurons. Our observation of robust relative threat signaling in male rats permits a direct test of this hypothesis in future studies.

The smallest and least expected population was High firing neurons signaling fear output and relative threat through firing decreases. High firing neurons are likely interneurons^58^ and may represent the animal’s current behavioral state or provide a readout of the current threat level. This information may be broadcasted within the VP and used by output neurons to construct signals for relative threat and salience. Intermediate firing neurons were the most selective, in that they showed relative threat signaling through firing decreases, but were not reward responsive. Intermediate firing neurons may be a proenkephalin (Penk) population specifically tuned to negative valence^59,^ but see a more recent study by Heinsbroek et.al^60^. In support, DREADD manipulation of VP Penk+ neurons has no effect on appetitive Pavlovian discrimination. Yet, DREADD excitation of VP Penk+ firing diminishes inhibitory avoidance, whereas DREADD inhibition has no impact^59^. Thus, exciting VP Penk neurons may oppose the normal reduction in firing to threat cues/contexts, disrupting threat behavior.

The patterned activity of Low firing neurons is comparable to studies showing VP populations with opposing changes in firing to rewarding and aversive cues. In one study, mice were trained to associate unique auditory cues with outcomes of differing valence (water vs. air puff) and size (small vs. large)^18^. The largest VP population showed firing increases to water cues that differentiated size (large > small), and subjects showed differential licking during cue presentation (large > small). These same neurons showed firing decreases to air puff cues that less clearly differentiated size (large ∼ small). However, behavior around air puff was not measured, so it is possible that the small and large aversive cues supported equivalent amounts of behavior. In the most recent study, monkeys were trained to associate visual cues with liquid reward, air puff or nothing (neutral)^40^. One VP population showed firing increases to the liquid reward cue, but firing decreases to the air puff cue. Yet, these same neurons showed comparable firing decreases to the neutral cue. Behaviorally, monkeys treated the neutral cue more similarly to the air puff cue. We observed firing decreases that differentiated threat cues associated with different foot shock probabilities (danger vs. uncertainty) and further differentiated threat cues from a neutral cue (safety). Differential decreases in VP firing may emerge in settings where threat probability estimates must be made and complete behavioral discrimination is observed.

Relative threat signaling through VP firing decreases is readily integrated into neural circuits permitting fine tuning of threat behavior. Low firing neurons are likely GABAergic output neurons^18,55,58^. Previous work has shown that suppressing VP activity promotes aversive behavior. So while optogenetic activation of all VP neurons/VP GABA neurons induces place preference^18,20,55^, inhibition of VP GABA neurons induces place aversion^55^. VP GABA firing decreases may simultaneously modulate VTA-driven reward behavior and BLA-driven threat behavior. VP GABA neurons directly project to dopamine neuron and GABA interneurons in the ventral tegmental area (VTA)^20,55,61^. VP GABA neurons also project to glutamate neuron and GABA interneurons in the BLA^21,62^. Consistent with a previous proposal^63^, threat-induced VP GABA firing decreases may increase VTA GABA activity, suppressing VTA dopamine firing to reduce reward behavior. At the same time, threat-induced VP GABA firing decreases may disinhibit BLA firing to promote threat behavior. By scaling firing decreases to degree of threat, VP neurons may permit fine modulation of VTA and BLA firing, thereby controlling the degree of threat response. Low/Intermediate firing neurons may also include cholinergic neurons, which densely project to the BLA^64,65^. Stimulating basal forebrain cholinergic terminals in the in BLA inhibits principal neurons that are modestly depolarized or at rest^66^. Thus, VP cholinergic and GABAergic firing decreases are positioned to suppress VTA dopamine firing and disinhibit BLA firing to precisely regulate threat behavior.

The VP is essential to reward behavior^15,23,28^ and recent evidence reveals the VP as a neural source of relative reward value^35^. A host of studies implicate the VP in aversive learning and behavior^18–20,36,39–41,55,63,67^. Here we reveal the VP as a neural source for relative threat. Detailing how VP relative threat signals shape firing and behavior through interactions with a larger neural circuit is likely to provide insight into the neural basis of adaptive and maladaptive threat behavior.

## Methods

### Experimental subjects

A total of 14 adult male Long Evans rats, weighing 250–275 g were obtained from Long Evans breeders maintained in the Boston College Animal Care Facility. The rats were single-housed on a 12 h light/dark cycle (lights on at 7:00 a.m.) with free access to water. Rats were maintained at 85% of their free-feeding body weight with standard laboratory chow (18% Protein Rodent Diet #2018, Harlan Teklad Global Diets, Madison, WI), except during surgery and post-surgery recovery. All protocols were approved by the Boston College Animal Care and Use Committee and all experiments were carried out in accordance with the NIH guidelines regarding the care and use of rats for experimental procedures.

### Electrode assembly

Microelectrodes consisted of a drivable bundle of sixteen 25.4 µm diameter Formvar-Insulated Nichrome wires (761500, A-M Systems, Carlsborg, WA) within a 27-gauge cannula (B000FN3M7K, Amazon Supply) and two 127 µm diameter PFA-coated, annealed strength stainless-steel ground wires (791400, A-M Systems, Carlsborg, WA). All wires were electrically connected to a nano-strip Omnetics connector (A79042-001, Omnetics Connector Corp., Minneapolis, MN) on a custom 24-contact, individually routed and gold immersed circuit board (San Francisco Circuits, San Mateo, CA). Sixteen individual recording wires were soldered to individual channels of an Omnetics connector. The sixteen wire bundle was integrated into a microdrive permitting advancement in ∼42 μm increments.

### Surgery

Stereotaxic surgery was performed aseptic conditions under isoflurane anesthesia (1-5% in oxygen). Carprofen (5 mg/kg, i.p.) and lactated ringer’s solution (10 mL, s.c.) were administered preoperatively. The skull was scoured in a crosshatch pattern with a scalpel blade to increase efficacy of implant adhesion. Six screws were installed in the skull to further stabilize the connection between the skull, electrode assembly and a protective head cap. A 1.4 mm diameter craniotomy was performed to remove a circular skull section centered on the implant site and the underlying dura was removed to expose the cortex. Nichrome recording wires were freshly cut with surgical scissors to extend ∼2.0 mm beyond the cannula. Just before implant, current was delivered to each recording wire in a saline bath, stripping each tip of its formvar insulation. Current was supplied by a 12 V lantern battery and each Omnetics connector contact was stimulated for 2 s using a lead. Machine grease was placed by the cannula and on the microdrive. For implantation dorsal to the VP, the electrode assembly was slowly advanced (∼100 μm/min) to the following coordinates: −0.08 mm form bregma, −2.05 mm lateral from midline, and −6.95 mm ventral from the cortex. Once in place, stripped ends of both ground wires were wrapped around two screws in order to ground the electrode. The microdrive base and a protective head cap were cemented on top of the skull using orthodontic resin (C 22-05-98, Pearson Dental Supply, Sylmar, CA), and the Omnetics connector was affixed to the head cap.

### Behavioral apparatus

All experiments were conducted in two, identical sound-attenuated enclosures that each housed a Pavlovian fear discrimination chamber with aluminum front and back walls retrofitted with clear plastic covers, clear acrylic sides and top, and a stainless steel grid floor. Each grid floor bar was electrically connected to an aversive shock generator (Med Associates, St. Albans, VT) through a grounding device. This permitted the floor to be grounded at all times except during shock delivery. An external food cup and a central nose poke opening, equipped with infrared photocells were present on one wall. Auditory stimuli were presented through two speakers mounted on the ceiling of enclosure. Behavior chambers were modified to allow for free movement of the electrophysiology cable during behavior; plastic funnels were epoxied to the top of the behavior chambers with the larger end facing down, and the tops of the chambers were cut to the opening of the funnel.

### Nose poke acquisition

Experimental procedure started with two days of pre-exposure in the home cage where rats received the pellets (Bio-Serv, Flemington, NJ) used for rewarded nose poking. Rats were then shaped to nose poke for pellet delivery in the behavior chamber using a fixed ratio schedule in which one nose poke yielded one pellet until they reached at least 50 nose pokes. Over the next 5 days, rats were placed on variable interval (VI) schedules in which nose pokes were reinforced on average every 30 s (VI-30, day 1), or 60 s (VI-60, days 2 through 5). For fear discrimination sessions, nose pokes were reinforced on a VI-60 schedule independent of auditory cue or foot shock presentation.

### Fear discrimination

Prior to surgery, each rat received eight 54-minutes Pavlovian fear discrimination sessions. Each session consisted of 16 trials, with a mean inter-trial interval of 3.5 min. Auditory cues were 10 s in duration and consisted of repeating motifs of a broadband click, phaser, or trumpet (listen or download: http://mcdannaldlab.org/resources/ardbark). Each cue was associated with a unique probability of foot shock (0.5 mA, 0.5 s): danger, *p*=1.00; uncertainty, *p*=0.25; and safety, *p*=0.00. Auditory identity was counterbalanced across rats. For danger and uncertainty-shock trials, foot shock was administered 2 s following the termination of the auditory cue. This was done in order to observe possible neural activity during the delay period is not driven by an explicit cue. A single session consisted of four danger trials, two uncertainty-shock trials, six uncertainty-omission trials, and four safety trials. The order of trial type presentation was randomly determined by the behavioral program, and differed for each rat, each session. After the eighth discrimination session, rats were given full food and implanted with drivable microelectrode bundles. Following surgical recovery, discrimination resumed with single unit recording. The microelectrode bundles were advanced in ∼42-84 μm steps every other day to record from new units during the following session.

### Single unit data acquisition

During recording sessions, a 1x amplifying headstage connected the Omnetics connector to the commutator via a shielded recording cable (Headstage: 40684-020 & Cable: 91809-017, Plexon Inc., Dallas TX). Analog neural activity was digitized and high-pass filtered via amplifier to remove low-frequency artifacts and sent to the Ominplex D acquisition system (Plexon Inc., Dallas TX). Behavioral events (cues, shocks, nose pokes) were controlled and recorded by a computer running Med Associates software. Timestamped events from Med Associates were sent to Ominplex D acquisition system via a dedicated interface module (DIG-716B). The result was a single file (.pl2) containing all time stamps for recording and behavior. Single units were sorted offline with a template-based spike-sorting algorithm (Offline Sorter V3, Plexon Inc., Dallas TX). Timestamped spikes and events (cues, shocks, nose pokes) were extracted and analyzed with statistical routines in Matlab (Natick, MA).

### Histology

Rats were deeply anesthetized using isoflurane and final electrode coordinates were marked by passing current from a 6 V battery through 4 of the 16 nichrome electrode wires. Rats were transcardially perfused with 0.9% biological saline and 4% paraformaldehyde in a 0.2 M Potassium Phosphate Buffered solution. Brains were extracted and post-fixed in a 10% neutral-buffered formalin solution for 24 h, stored in 10% sucrose/formalin, frozen at −80°C and sectioned via sliding microtome. In order to identify VP boundaries, we performed immunohistochemistry for substance P (primary antibody, rabbit anti-substance P, Immunostar, Hudson, WI; secondary antibody, Alexa Fluor 594 donkey anti-rabbit, Jackson ImmunoResearch Laboratories, West Grove, PA), and NeuroTrace™(1:200, Thermo Fisher Scientific, Waltham, MA). Sections were mounted on coated glass slides, coverslipped with Vectashield mounting medium without DAPI (Vector Laboratories, Burlingame, CA), and imaged using a fluorescent microscope (Axio Imager Z2, Zeiss, Thornwood, NY). Electrode placements were reconstructed by subtracting the distance driven between recording sessions from the final recording site. All the recording sites within the boundaries of VP were included in analyses^68^.

### Statistical analysis

### 95% bootstrap confidence interval

95% bootstrap confidence intervals were constructed for suppression ratios, differential firing, and beta coefficients using the bootci function in Matlab. For each bootstrap, a distribution was created by sampling the data 1,000 times with replacement. Studentized confidence intervals were constructed with the final outputs being the mean, lower bound and upper bound of the 95% bootstrap confidence interval. Suppression ratios, differential firing, and beta coefficients were said to be observed when the 95% confidence interval did not include zero.

### Calculating suppression ratios

Fear was measured by suppression of rewarded nose poking, calculated as a ratio: [(baseline poke rate - cue poke rate) / (baseline poke rate + cue poke rate)]. The baseline nose poke rate was taken from the 20 s prior to cue onset and the cue poke rate from the 10 s cue period. Suppression ratios were calculated for each trial using only that trial’s baseline. A ratio of ‘1’ indicated high fear, ‘0’ low fear, and gradations between intermediate levels of fear. Suppression ratios were analyzed using ANOVA with cue (danger, uncertainty, and safety) as a factor (Fig. 1f). Uncertainty-shock and uncertainty-omission trials were collapsed because they did not differ for suppression ratio; during cue presentation, rats did not know the current uncertainty trial type. F statistic, *p* value, partial eta squared (η_p_^2^) and observed power (op) are reported for significant main effects and interactions. The distribution of suppression ratios was visualized using the plotSpread function for Matlab (https://www.mathworks.com/matlabcentral/fileexchange/37105-plot-spread-points-beeswarm-plot).

### Identifying cue-responsive neurons

Single units were screened for cue responsiveness by comparing raw firing rate (spikes/s) during the 10 s baseline period just prior to cue onset to firing rate during the first 1 s and last 5 s of danger, uncertainty and safety using a paired, two-tailed t-test (*p*<0.05). A neuron was considered cue-responsive if it showed a significant increase or decrease in firing to any cue in either period. A full Bonferroni correction (0.5/6) was not performed because this criterion was too stringent, resulting in many cue-responsive neurons being omitted from analysis.

### Firing and waveform characteristics

The following characteristics were determined for each cue-responsive neuron: baseline firing rate, coefficient of variance, coefficient of skewness, waveform half-duration, and waveform amplitude ratio (Fig.S1). Baseline firing rate was mean firing rate (spikes/s) during the 10 s baseline period just prior to cue onset. Coefficient of variance was calculated by 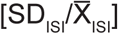, in which SD_ISI_ was the standard deviation of inter-spike interval, and 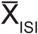 was the mean inter-spike interval. Coefficient of variance is a relative measure of the variability of spike firing, with small values indicating less variation in inter-spike intervals (more regular firing), and large values more variability (less regular firing)^47,48^. Coefficient of skewness was calculated by 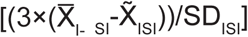, in which 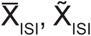, and SD_ISI_ were the mean, median and standard deviation of inter-spike interval, respectively. Coefficient of skewness is a measure of the asymmetry of the distribution of the inter-spike intervals, with positive values indicating longer intervals (less regular firing) and negative values indicating shorter intervals (more regular firing)^48^. Waveform amplitude ratio was calculated by [(N-P)/(N+P)], in which P was the y-axis distance between the initial value and peak initial hyperpolarization, and N was the y-axis distance between the peak initial value and valley of depolarization. Values near zero indicate a relatively large initial hyper-polarization while values near one indicate a relatively small initial hyperpolarization^49,50^. Waveform half-duration was calculated by [D/2)], in which D was the x-axis distance between the valley of depolarization and the peak of after-hyperpolarization and smaller values indicate narrower waveforms^49,50^.

### K-means clustering

We used k-means clustering to identify cue-inhibited subpopulations. Clustering was performed using the Matlab kmeans function. K-means clustering used baseline firing and four additional characteristics (coefficient of variance, coefficient of skewness, waveform half-duration, and waveform amplitude ratio) and identified three clusters within the population.

### Z-score normalization

For each neuron, and for each trial type, firing rate (spikes/s) was calculated in 250 ms bins from 20 s prior to cue onset to 20 s following cue offset, for a total of 200 bins. Mean firing rate over the 200 bins was calculated by averaging all trials for each trial type. Mean differential firing was calculated for each of the 200 bins by subtracting mean baseline firing rate (10 s prior to cue onset), specific to that trial type, from each bin. Mean differential firing was Z-score normalized across all trial types within a single neuron, such that mean firing = 0, and standard deviation in firing = 1. Z-score normalization was applied to firing across the entirety of the recording epoch, as opposed to only the baseline period, in case neurons showed little/ no baseline activity. As a result, periods of phasic, excitatory and inhibitory firing contributed to normalized mean firing rate (0). For this reason, Z-score normalized baseline activity can differ from zero. Z-score normalized firing was analyzed with ANOVA using cue, and bin as factors. F and *p* values are reported, as well as partial eta squared (η _p_^2^) and observed power (op). For reward firing, firing rate (spikes/s) was calculated in 250 ms bins from 2 s prior to reward delivery to 2 s following reward delivery, for a total of 16 bins. Mean differential firing was calculated for each of the 16 bins by subtracting pre-reward firing rate (mean of 1 s prior to reward delivery).

### Heat plot and color maps

Heat plots were constructed from normalized firing rate using the imagesc function in Matlab (Fig. 2). Perceptually uniform color maps were used to prevent visual distortion of the data^69^.

### Population and single unit firing analyses

Population cue firing was analyzed using ANOVA with cue (danger, uncertainty, and safety) and bin (250 ms bins from 2 s prior to cue onset to 2 s following cue offset) as factors (Fig. 3 and Fig. 6a, b). Uncertainty trial types were collapsed because they did not differ firing analysis. This was expected, during cue presentation rats did not know the current uncertainty trial type. F statistic, *p* value, partial eta squared (η_p_^2^) and observed power (op) are reported for main effects and interactions. The 95% bootstrap confidence intervals were reconstructed for normalized firing to each cue (compared to zero), as well as for differential firing (danger vs. uncertainty) and (uncertainty vs. safety), during cue onset (first 1 s cue interval), late cue (last 5 s cue interval), and delay periods (2 s following cue offset). The distribution of single unit firing was visualized using a plotSpread function for Matlab. Population reward firing was analyzed using repeated measures ANOVA with bin (250 ms bins from 2 s prior to reward delivery to 2 s following reward delivery) as factor (Fig. 5 a-d and Fig. 6e, f). The 95% bootstrap confidence intervals were reconstructed for normalized firing to reward during pre (500 ms prior to reward delivery), and post (first 500 ms following reward delivery) (compared to zero), as well as for differential firing (pre vs. post). Relationships between cue firing (danger vs. uncertainty, and uncertainty vs. safety; Fig. S2 and Fig. S4), as well as between reward and cue firing (Fig. 5-e-m Fig. 6g) were determined by calculating the R^2^ and *p* value for the Pearson’s correlation coefficient. Population firing was analyzed using repeated measures ANOVA with bin (250 ms bins from 2 s prior to nose poke cessation to 2 s following nose poke cessation) as factor (Fig. S3 and Fig. S5). The 95% bootstrap confidence intervals were reconstructed for normalized firing during pre (500 ms prior to nose poke cessation), and post (first 500 ms following nose poke cessation) (compared to zero), as well as for differential firing (pre vs. post).

### Single unit, linear regression

Single unit, linear regression was used to determine the degree to which fear output and/or relative threat explained trial-by-trial variation in firing of single neurons in a specific time interval. For each regression, all 16 trials from a single session were ordered by type. Z-score normalized firing rate was specified for the interval of interest. The relative threat regressor was the foot shock probability associated with the specific cue (danger = 1.00, uncertainty = 0.25, safety = 0.00). The fear output regressor was the suppression ratio for the entire cue, for that specific trial. Regression (using the regress function in Matlab) required a separate, constant input. The regression output was the beta coefficient for each regressor (relative threat and fear output), quantifying the strength (greater distance from zero = stronger) and direction (>0 = positive) of the predictive relationship between each regressor and single unit firing. ANOVA was used to analyze beta coefficients, exactly as described for normalized firing rate (Fig. 4 and Fig. 6c, d). The 95% bootstrap confidence intervals were reconstructed for beta coefficients (compared to zero), as well as for relative threat vs. fear output during cue (10 s cue interval), and delay (2 s following cue offset) periods. The distribution of single unit beta coefficients visualized using a plotSpread function for Matlab.

### Data and software availability

Full electrophysiology data set will be uploaded to http://crcns.org/ upon acceptance for publication.

### Additional resources

Med Associates programs used for behavior and Matlab programs used for behavioral analyses are made freely available at our lab website: http://mcdannaldlab.org/resources

## Acknowledgements

We thank Bret Judson and the Boston College Imaging Core for infrastructure and support.

