## Supplemental Files for "Ventral pallidum neurons signal relative threat"

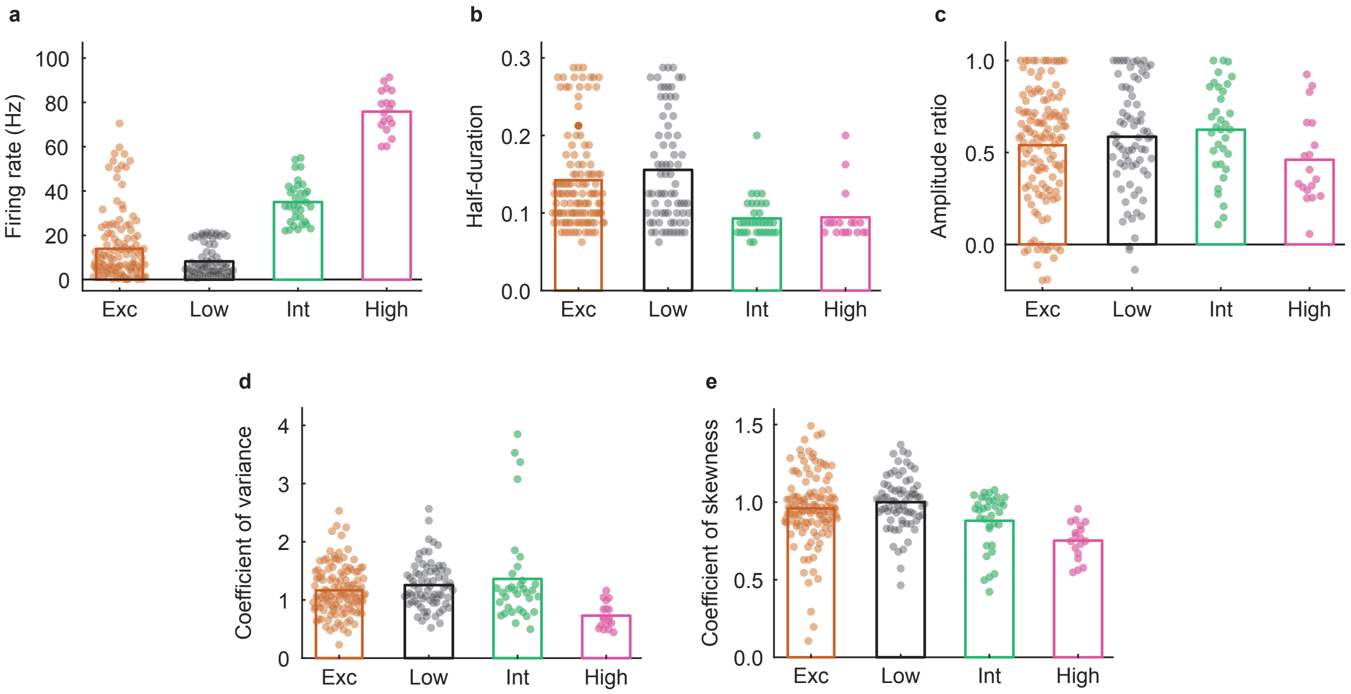


**Fig. S1. Firing and waveform characteristics of cue-excited and cue-inhibited neurons.** Mean (bar) and individual (data points) (**a**) firing rate (**b**) waveform half-duration (**c**) waveform amplitude ratio (**d**) coefficient of variance, and (**e**) coefficient of skewness during a 10 s baseline period just prior to cue onset for cue-excited (Exc, n = 131, maroon), Low firing (Low, n = 74, black), Intermediate firing (Int, n = 34, turquoise), and High firing (High, n = 18, pink) neurons.

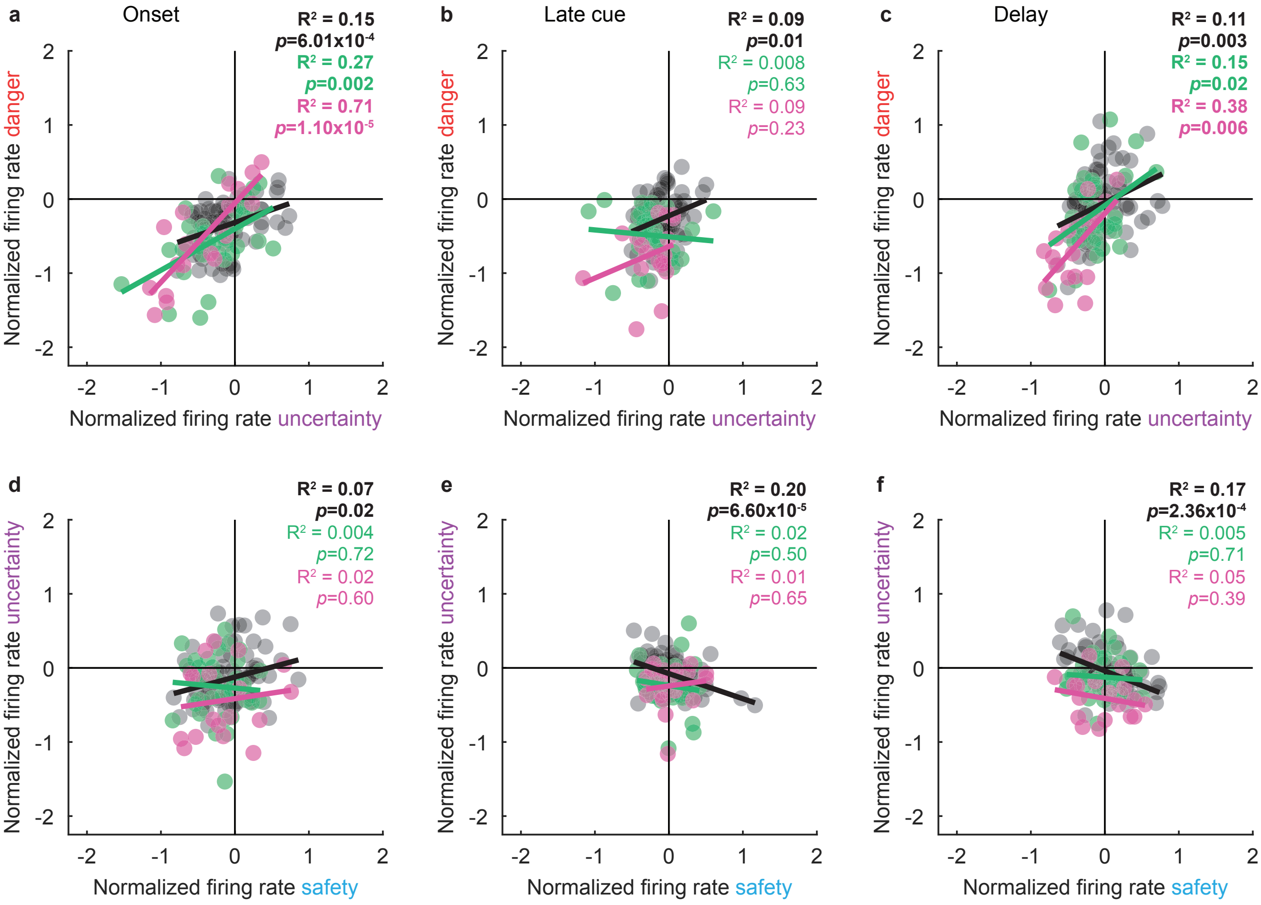


**Fig. S2. Cue firing relationships in cue-inhibited neurons.** Scatterplots for mean normalized firing rate to danger (red) vs. uncertainty (purple) are shown for (**a**, onset) the first 1 s cue interval, (**b**, late cue) last 5 s cue interval, and (**c**, delay) 2 s following cue offset for the cue-inhibited population (Low firing, n = 74, black; Intermediate firing, n = 34, turquoise; High firing, n = 18, pink). Trendline, the square of the Pearson correlation coefficient ($R^2$) and associated p value (*p*) are shown for each cluster. (**d-f**) Scatterplots for mean normalized firing rate to uncertainty (purple) vs. safety (blue) shown, as in a-c.

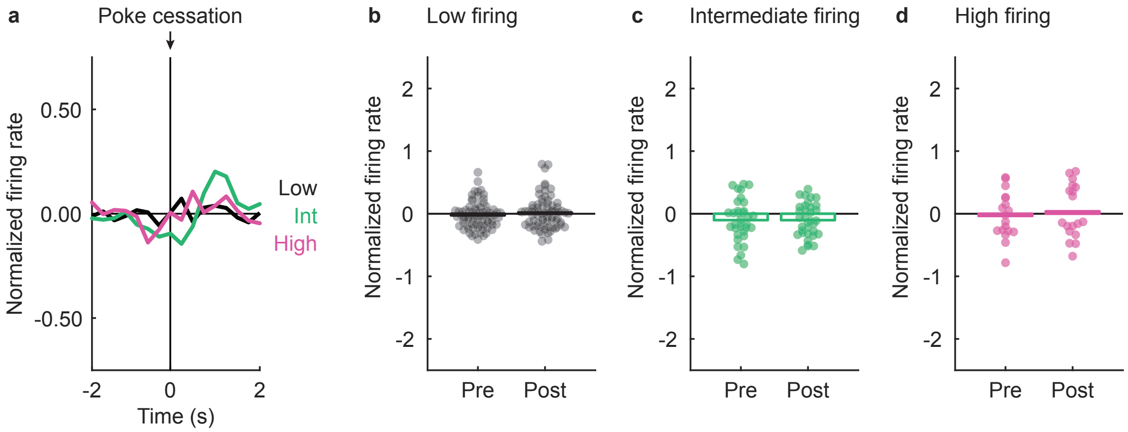


**Fig. S3. Nose poke cessation is insufficient to drive activity of cue-inhibited neurons.** (**a**) Mean normalized firing rate is shown 2 s prior to and 2 s after nose poke cessation for the Low (Low, n = 74, black), Intermediate (Int, n = 34, turquoise), and High (High, n = 18, pink) firing neurons. All cue-inhibited neurons were unresponsive to nose poke cessation. Nose poke cessation is indicated by black arrow. (**b-d**) Mean (bar) and individual (data points), normalized firing rate for (**b**) Low (n = 74, black), (**c**) Intermediate (n = 34, turquoise), and (**d**) High (n = 18, pink) firing neurons are shown during 500 ms interval prior (pre) to and 500 ms interval after (post) nose poke cessation.

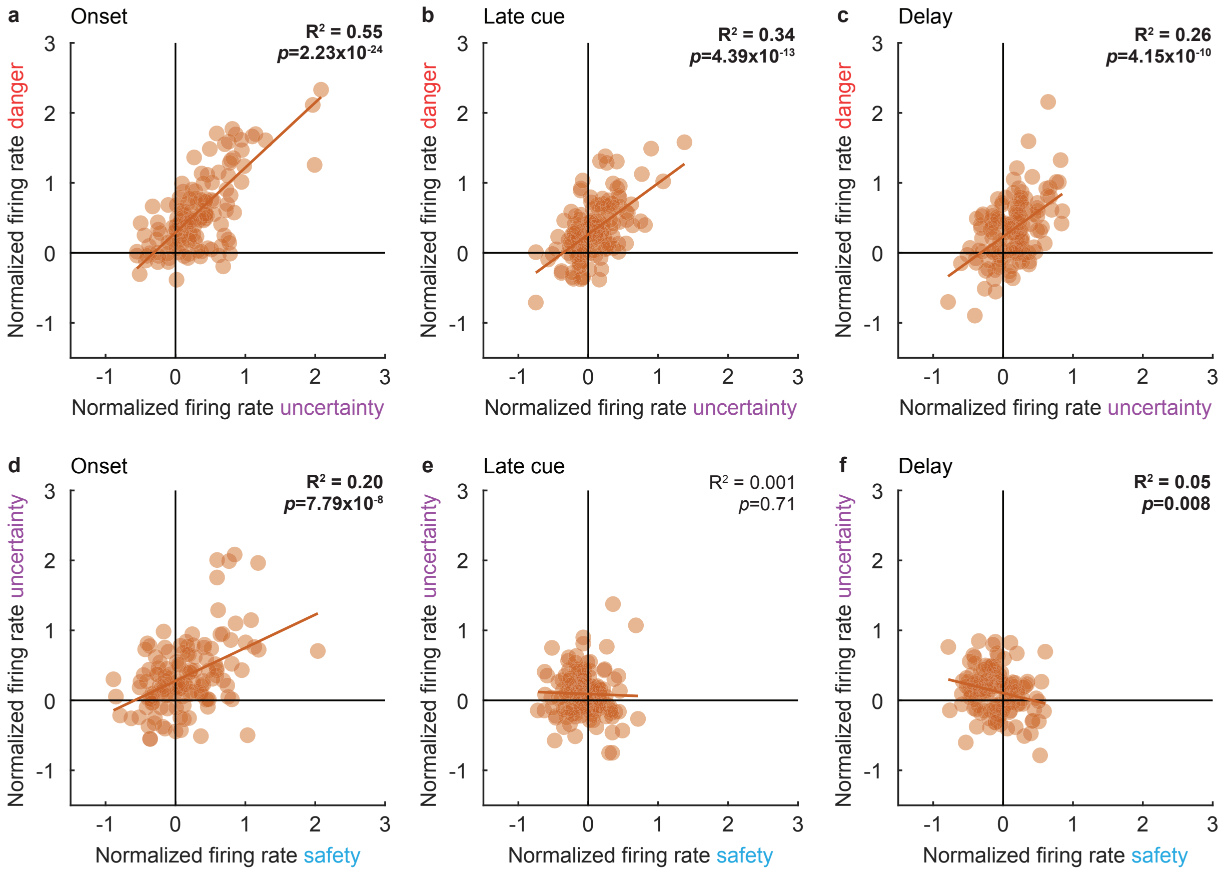


**Fig. S4. Cue firing relationships in cue-excited neurons.** Scatterplots for mean normalized firing rate to danger (red) vs. uncertainty (purple) are shown for (**a,** onset) the first 1 s cue interval, (**b,** late cue) last 5 s cue interval, and (**c,** delay) 2 s following cue offset for the cue-excited population (n = 131, maroon). Trendline, the square of the Pearson correlation coefficient ($R^2$) and associated p value (*p*) are shown for each cluster. (**d-f**) Scatterplots for mean normalized firing rate to uncertainty (purple) vs. safety (blue) shown, as in a-c.

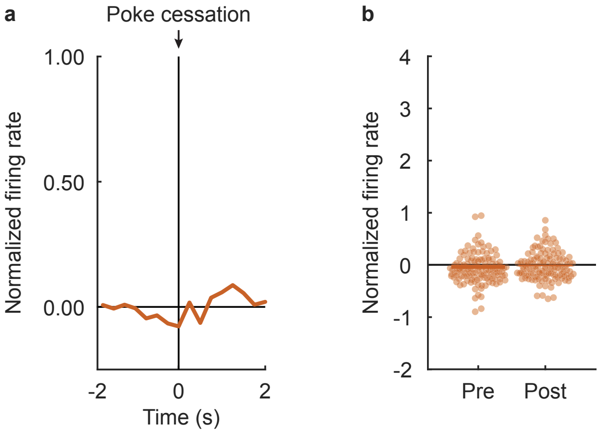


**Fig. S5. Nose poke cessation is insufficient to drive activity of cue-excited neurons.** (**a**) Mean normalized firing rate is shown 2 s prior to and 2 s after nose poke cessation for cue-excited neurons (n = 131, maroon). Nose poke cessation is indicated by black arrow. (**b**) Mean (bar) and individual (data points), normalized firing rate for cue-excited neurons (n = 131, maroon) are shown during 500 ms interval prior (pre) to and 500 ms interval after (post) nose poke cessation.
